# Hierarchical organization of the human cell from a cancer coessentiality network

**DOI:** 10.1101/328880

**Authors:** Eiru Kim, Merve Dede, Walter F. Lenoir, Gang Wang, Sanjana Srinivasan, Medina Colic, Traver Hart

## Abstract

Genetic interactions mediate the emergence of phenotype from genotype. Systematic survey of genetic interactions in yeast showed that genes operating in the same biological process have highly correlated genetic interaction profiles, and this observation has been exploited to infer gene function in model organisms. Systematic surveys of digenic perturbations in human cells are also highly informative, but are not scalable, even with CRISPR-mediated methods. As an alternative, we developed an indirect method of deriving functional interactions. We show that genes having correlated knockout fitness profiles across diverse, non-isogenic cell lines are analogous to genes having correlated genetic interaction profiles across isogenic query strains, and similarly implies shared biological function. We constructed a network of genes with correlated fitness profiles across 400 CRISPR knockout screens in cancer cell lines into a “coessentiality network,” with up to 500-fold enrichment for co-functional gene pairs, enabling strong inference of human gene function. Modules in the network are connected in a layered web that gives insight into the hierarchical organization of the cell.

## Introduction

Genetic interactions govern the translation of genotype to phenotype at every level, from the function of subcellular molecular machines to the emergence of complex organismal traits. In the budding yeast *Saccharomyces cerevisiae*, systematic genetic deletion studies showed that only ∼1,100 of its ∼6,000 genes (∼20%) were required for growth under laboratory conditions (Giaever et al., 2002). A systematic survey of digenic knockouts, however, yielded hundreds of thousands of gene pairs whose double knockout induced a fitness phenotype significantly more severe (synergistic genetic interactions) or less severe (suppressor interactions) than expected from each gene’s single mutant fitness (Costanzo et al., 2010, 2016; Tong et al., 2001), with triple-mutant screens adding yet another layer of complexity (Kuzmin et al., 2018). When trying to decipher the genetic contribution to as simple a phenotype as fitness, then, there are vastly more candidate explanations involving genetic interactions than monogenic fitness effects. Moreover, the impact of each gene variant not only depends on the sum of all other genetic variants in the cell, but also is strongly influenced by the cell’s environment (Bandyopadhyay et al., 2010; Hillenmeyer et al., 2008).

Patterns of genetic interaction are deeply informative. Genetic interactions frequently occur either within members of the same pathway or process (“within pathway interactions”) or between members of parallel pathways (“between pathway interactions”) (Kelley and Ideker, 2005). When assayed systematically, the result is that genes that operate in the same biological process tend to interact genetically with the same sets of other genes in discrete, related pathways, culminating in highly correlated genetic interaction profiles across a diverse panel of genetic backgrounds or “query strains.” This observation has been exploited extensively to infer gene function in model organisms and, on a smaller scale, in human cells based on similarity of genetic interaction profiles (Lehner et al., 2006; Horn et al., 2011; Bassik et al., 2013; Kampmann et al., 2013; Roguev et al., 2013; Costanzo et al., 2016). Therefore, beyond the specific interactions themselves, a gene’s pattern of fitness phenotypes across a diverse set of backgrounds can inform our knowledge of that gene’s function.

Translating these concepts into human cells has proved biologically and technically challenging. The *S. cerevisiae* genome has less than one-third the number of protein coding genes as humans, and, despite the quantum leap in technology that the CRISPR/Cas system offers to mammalian forward genetics, yeast remains far simpler to perturb reliably in the lab. Several groups have applied digenic perturbation technologies, using both shRNA and CRISPR, to find cancer genotype-specific synthetic lethals for drug targeting (Han et al., 2017; Najm et al., 2018; Sage et al., 2017; Shen et al., 2017; Wong et al., 2016); (Du et al., 2017) and to identify genetic interactions that enhance or suppress phenotypes related to drug and toxin resistance (Bassik et al., 2013; Jost et al., 2017; Roguev et al., 2013). The current state of the art in CRISPR-mediated gene perturbation relies on observations from three independent guide RNA (gRNA) targeting each gene, or nine pairwise perturbations for each gene pair, plus non-targeting or other negative controls. The largest such mapping to date puts the scale of the problem in stark terms: Han *et al.* use a library of 490,000 gRNA doublets – seven times larger than a latest generation whole-genome, single-gene knockout library – to query all pairs of 207 target genes, or ∼0.01% of all gene pairs in the human genome (Han et al., 2017).

An additional dimension of the scale problem is that of backgrounds. Whereas one strain of yeast was systematically assayed in fixed media and environmental conditions to create a reference genetic interaction network, no such reference cell exists for humans. Indeed first-generation whole-genome CRISPR screens in cancer cell lines demonstrated that one of the features associated with the hugely increased sensitivity of CRISPR over shRNA (Hart et al., 2014, 2015) was the ability to resolve tissue- and genetic-driven differences in gene essentiality, as well as the unexpected variation in gene essentiality in cell lines with ostensibly similar genetic backgrounds (Hart et al., 2015; Wang et al., 2014).

Nevertheless, small-scale, targeted genetic interaction screens in human cells using both shRNA and CRISPR showed that the architecture of the genetic interaction network holds true across species. Positive and negative genetic interactions within pathways and between related biological processes yield a correlation network with the same properties: genes with similar profiles of genetic interaction across different backgrounds are often in the same process or complex, providing a strong basis for inference of gene function (Bassik et al., 2013, 2013; Horn et al., 2011; Kampmann et al., 2013, 2014; Roguev et al., 2013). Since digenic perturbation screens are difficult to scale, we considered whether indirect methods of determining functional genomic information might be effective. Whole-genome CRISPR knockout screens have been performed in over 400 cancer and immortalized cell lines, with the bulk coming from Project Achilles using standardized protocols and reagents (Aguirre et al., 2016; Meyers et al., 2017; Tsherniak et al., 2017). We hypothesized that genes having correlated knockout fitness profiles across diverse cell lines would be analogous genes having correlated genetic interaction profiles across specified query backgrounds in the same cells, and would similarly imply shared biological function. We constructed a network of genes with correlated essentiality scores into a “coessentiality network,” from which we identified clusters of genes with high functional coherence. The network provides powerful insight into functional genomics, cancer targeting, and the capabilities and limitations of CRISPR-mediated genetic screening in human cell lines.

## Results & Discussion

We considered CRISPR and shRNA whole-genome screen data from multiple libraries and laboratories (Avana (Doench et al., 2014; Meyers et al., 2017), GeCKOv2 (Aguirre et al., 2016), TKO (Hart et al., 2015, 2017; Steinhart et al., 2017), Sabatini (Wang et al., 2014, 2017) the Moffat shRNA library (Koh et al., 2012; Marcotte et al., 2012, 2016; Medrano et al., 2017)) and other large data sets (McDonald et al., 2017; Tsherniak et al., 2017) (Figure 1a and Supplementary Table 1). From raw read count data, we used the BAGEL pipeline (described in (Hart and Moffat, 2016) and improved here; see Supplementary Methods) to generate Bayes Factors for each gene in each cell line. We removed nontargeting and nonhuman gene controls and quantile normalized each data set, yielding an essentiality score where a positive value indicates a strong knockout fitness defect and a negative value generally implies no phenotype (see Supplemental Methods for details). Each gene therefore has an “essentiality profile” of its scores across the screens in that data set.

**Figure 1.**
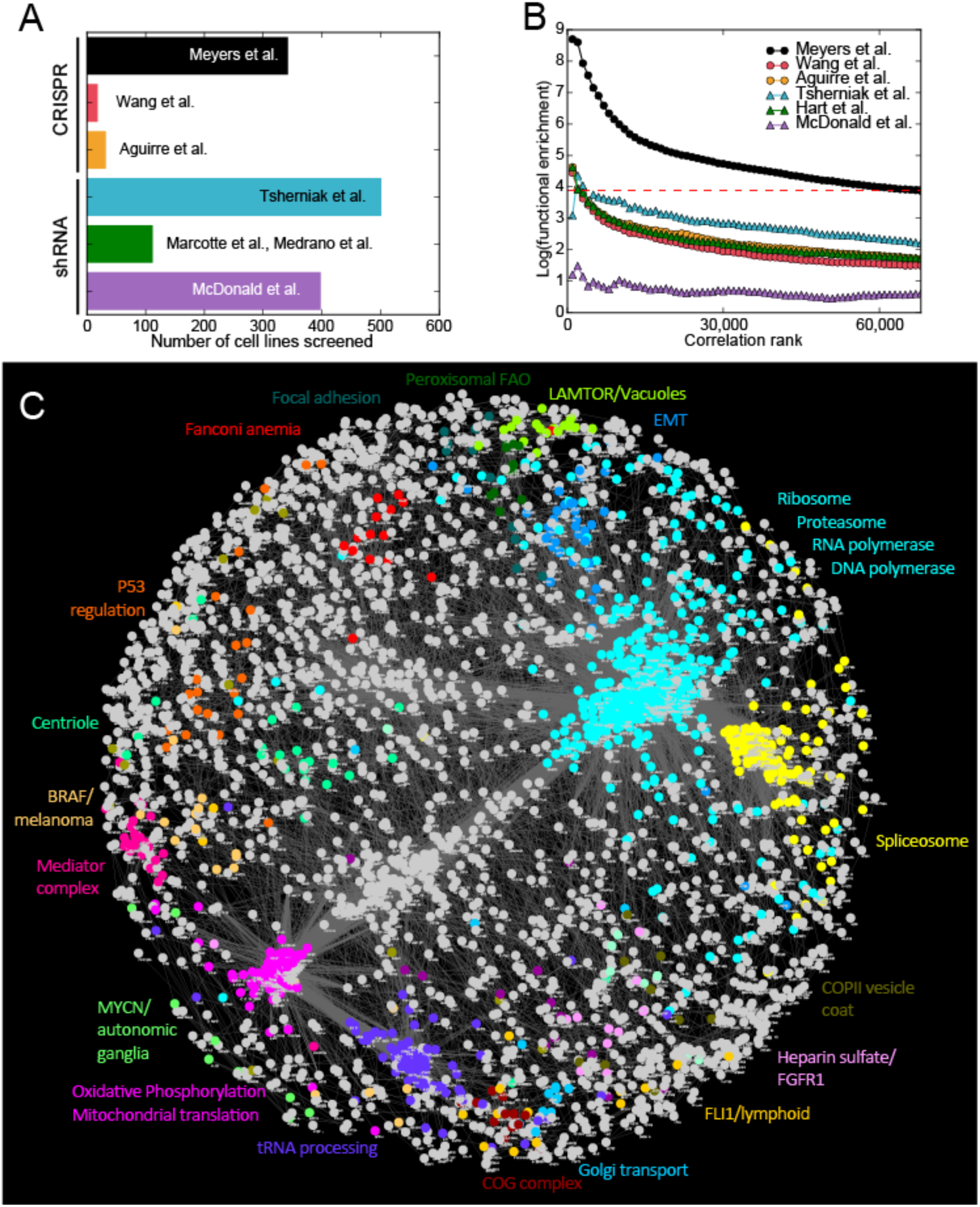
The coessentiality network. (A) CRISPR and shRNA screens analyzed for this study. (B) Measuring functional enrichment. For each data set, pairwise correlations of knockout/knockdown fitness profiles were ranked, binned (n=1000), and measured for enrichment for shared KEGG terms. Data from Meyers et al (“Avana data”) carries significantly more functional information than other data sets. (C) The Cancer Coessentiality Network, derived from Avana data, contains 3,483 genes connected by 68,813 edges. Selected modules, derived by an unbiased clustering algorithm and color-coded, demonstrate the functional coherence of the network.

For each data set, we ranked gene pairs by correlated essentiality profiles and measured the enrichment for co-functional pairs (see Methods). Data from Meyers et al, where CRISPR knockout screens were conducted using the Avana library in 342 cancer cell lines, showed the strongest enrichment for co-functional gene pairs (Figure 1b), likely due to the relatively high quality of the screens (Supplementary Figure 1) as well as the lineage and genetic diversity of the cells being screened. In contrast, screens from Wang et al. (Wang et al., 2014, 2017) were equally high quality but were performed only in 17 AML cell lines with correspondingly limited diversity. To further increase the co-functionality signal, we removed screens with poor performance and only considered genes that were hits in at least 3 of the remaining screens; filtering resulted in an additional twofold enrichment for co-functional gene pairs (Figure 1b and Supplementary Figure 1). The filtered data from Meyers et al. (Meyers et al., 2017) (n=276 cell lines; 5,387 genes; hereafter “Avana data”) was used for all subsequent analysis. We selected gene pairs with a Bonferroni-corrected P-value < 0.05 and combined them into a network, the Cancer Coessentiality Network, containing 3,327 genes connected by 68,641 edges (Figure 1c). The network is highly modular, with clusters showing strong functional coherence, similar to the networks directly inferred from correlated yeast genetic interaction profiles.(Costanzo et al., 2010, 2016)

### Essential genes specific to oncogenic contexts

The data underlying the Cancer Coessentiality Network is derived from well-characterized cancer cell lines from 30+ lineages, representing the major oncogenic mutation profiles common to those cancers. Many clusters in the network can therefore be associated with specific tissues and cancer-relevant genotypes. By testing cluster-level essentiality profiles for tissue specificity (see Supplementary Methods), we identified a number of clusters that correspond to tissue specific cancers (Figure 2a), which in turn contain the characteristic oncogenes. For example, Cluster 14 (Figure 2b) consists of BRAF and related genes that are highly specific to BRAF-mutated melanoma cells (P<10^−12^; Figure 2c). The cluster contains other elements of the MAP kinase pathway (MAP2K1, MAPK1, DUSP4), indicating their essentiality in BRAF-mutant cells, supporting efforts to incorporate ERK inhibitors into combinatorial therapies to overcome resistance to targeted BRAF treatments (Smalley and Smalley, 2018). This example highlights the utility of this indirect approach to identify synthetic lethal interactions: genes co-essential with oncogenes are synthetic lethals. Beyond the downstream elements of the MAPK pathway itself, the BRAF/melanoma cluster also contains the transcription factors (TFs) Melanogenesis Associated Transcription Factor (MITF); two developmental Sry-related box (Sox) genes, SOX9 and SOX10; and mesenchymal marker ZEB2, indicative of the non-epithelial origin of melanocyte cells and providing insight into the genetic requirements for tissue differentiation in this lineage.

**Figure 2.**
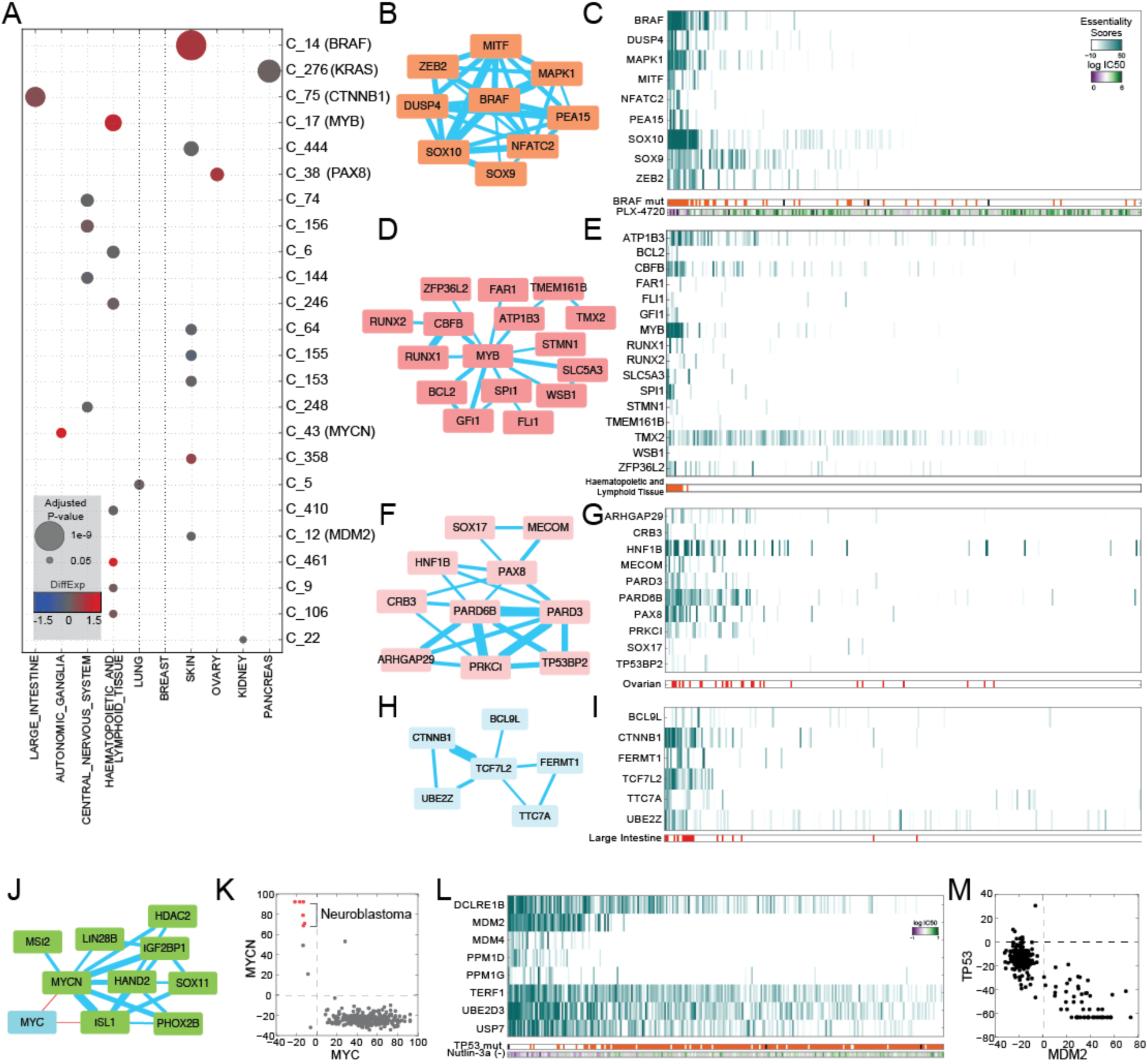
Cancer-specific features of the network. (A) Clusters of genes were evaluated for tissue specificity (size of circles) and differential mRNA expression of genes in the cluster (color of circles). (B) Cluster 14 (BRAF cluster); nodes are genes in cluster and edges reflect strength of correlation of fitness profile. (C) Heatmap of essentiality profiles of genes in BRAF cluster, ranked by median essentiality score. Gene essentiality in the cluster is associated with PBRAF mutation (P<10^−23^) and sensitivity to BRAF inhibitor PLX-4720 (P<10^−7^). (D,E) Network and heatmap of MYB-related cluster. (F,G) PAX8-associated cluster. (H,I) B-catenin cluster. (J) MYCN neuroblastoma cluster is anticorrelated with MYC.(K) MYC, MYCN essentiality is mutually exclusive. (L) MDM2 cluster heatmap is associated with TP53 mutation status (P<10^−13^) and sensitivity to Nutlin-3a (P<10^−14^). (M) MDM2 vs TP53 essentiality. TP53 essentiality scores < −50 indicate tumor suppressor role.

Similar observations hold for other tissue-specific oncogenes. Cluster 17, essential in lymphoid cell lines (P<10^−7^), contains oncogene FLI1 and tissue-specific transcription factor MYB (Figure 2d-e), and cluster 38 is enriched for ovarian cancer cells (P<10^−7^) and carries lineage-specific TF PAX8, previously shown to be essential in these cells (Cheung et al., 2011) (Figure 2f-g). Cluster 75, essential in colorectal cancer cells (P<10^−9^), contains β-catenin (CTNNB1) and transcription factor partner TCF7L2 (Figure 2h-i); both are linked to E2 ubiquitin ligase UBE2Z, which mediates UBA6-specific suppression of EMT (Liu et al., 2017), indicating a functional linkage with β-catenin signaling. Additional tissue- and oncogene-driven clusters delineating breast cancer subtypes, joint cyclin/cyclin-dependent kinase (CDK) dependencies, candidate KRAS synthetic lethals, and glioblastoma-specific essential genes are shown in Supplementary Figure 2.

Neuroblastoma cells require MYCN, the neuroblastoma-specific paralog of the MYC oncogene (Huang and Weiss, 2013), as well as nervous system developmental transcription factor SOX11 (Potzner et al., 2010) (Figure 2j). Interestingly, MYC is highly essential in virtually all non-neuroblastoma cell lines, resulting in a relatively uncommon anticorrelation in essentiality profiles (r=-0.49; P<10^−17^; Figure 2k). While this negative correlation is driven by mutual exclusivity in tissue essentiality, we also observe anticorrelation in tumor suppressors and their repressors in the same cells. CRISPR knockout of tumor suppressors in cells carrying wildtype alleles frequently results in increased growth rate, which manifests as extreme negative BF scores with our updated BAGEL algorithm (see Supplementary Methods). TP53 has these extreme negative values in melanoma cells and others with unmutated genes, resulting in strong anticorrelation with TP53 suppressors MDM2 (r=-0.86, P<10^−81^), MDM4 (r=-0.61, P<10^−28^), and PPM1D (r=-0.72, P<10^−44^) (Figure 2l-m).

While TP53 shows the characteristic extreme negative essentiality score of a tumor suppressor gene in wildtype backgrounds, surprisingly, it causes a growth defect when knocked out in three cell lines: HCT1143 breast cancer, PC14 lung cancer, and NB4 AML cells (Supplementary Figure 3a). All three carry the R248Q oncogenic mutation; in fact, R248Q is weakly predictive of TP53 essentiality generally, and strongly predictive when it is the only P53 mutation detected (Supplementary Figure 3b). Nor is this the only case where a tumor suppressor in one background is an essential gene in another: the von Hippel-Landau tumor suppressor gene VHL shows no phenotype in renal cancer cells, where the gene is nearly universally deleted, but is essential specifically in BTFC-909 renal carcinoma cells which lack the characteristic Chr3 copy loss (Sinha et al., 2017). In contrast, VHL shows a fitness defect when knockout out in most other backgrounds (Supplementary Figure 3c). The essentiality profile for VHL is strongly correlated with EGLN1 (commonly called PHD2), an oxygen sensor that hydroxylates hypoxia response genes HIF1A and HIF2A, marking them for degradation by the VHL complex in normoxic environments (Berra et al., 2003). EGLN1 essentiality is overrepresented in melanoma cells (P<10^−4^, rank-sum test; essential in 14 of 22 skin cancer cell lines).

### A high-precision functional interaction map of human genes

These examples indicate the breadth and precision of the coessentiality network, but represent results from hypothesis-guided queries. In an effort to learn novel associations from the data, we tested each cluster for its correlation with cell lineage as well as correlation with gene expression, mutation, and copy number amplification of all genes both inside and outside the cluster to identify underlying molecular genetic drivers of modular, emergent essentiality. We identified 270 genes in 30 clusters whose essentiality profiles strongly correlated with their own copy number profiles but not their expression profiles (Figure 3a). As copy number amplification is a known source of false positives in CRISPR screens, we labeled these clusters as amplification artifacts. An additional 56 genes in 11 clusters showed significant association with both copy number and expression profiles. These clusters notably include KRAS amplifications in pancreatic and colorectal cancer (Cluster 276), ERBB2 amplifications in breast and other cancers (Cluster 52), and CCNE1 overexpression/RB1 mutation. (Cluster 101), consistent with well-studied patterns of oncogenesis.

**Figure 3.**
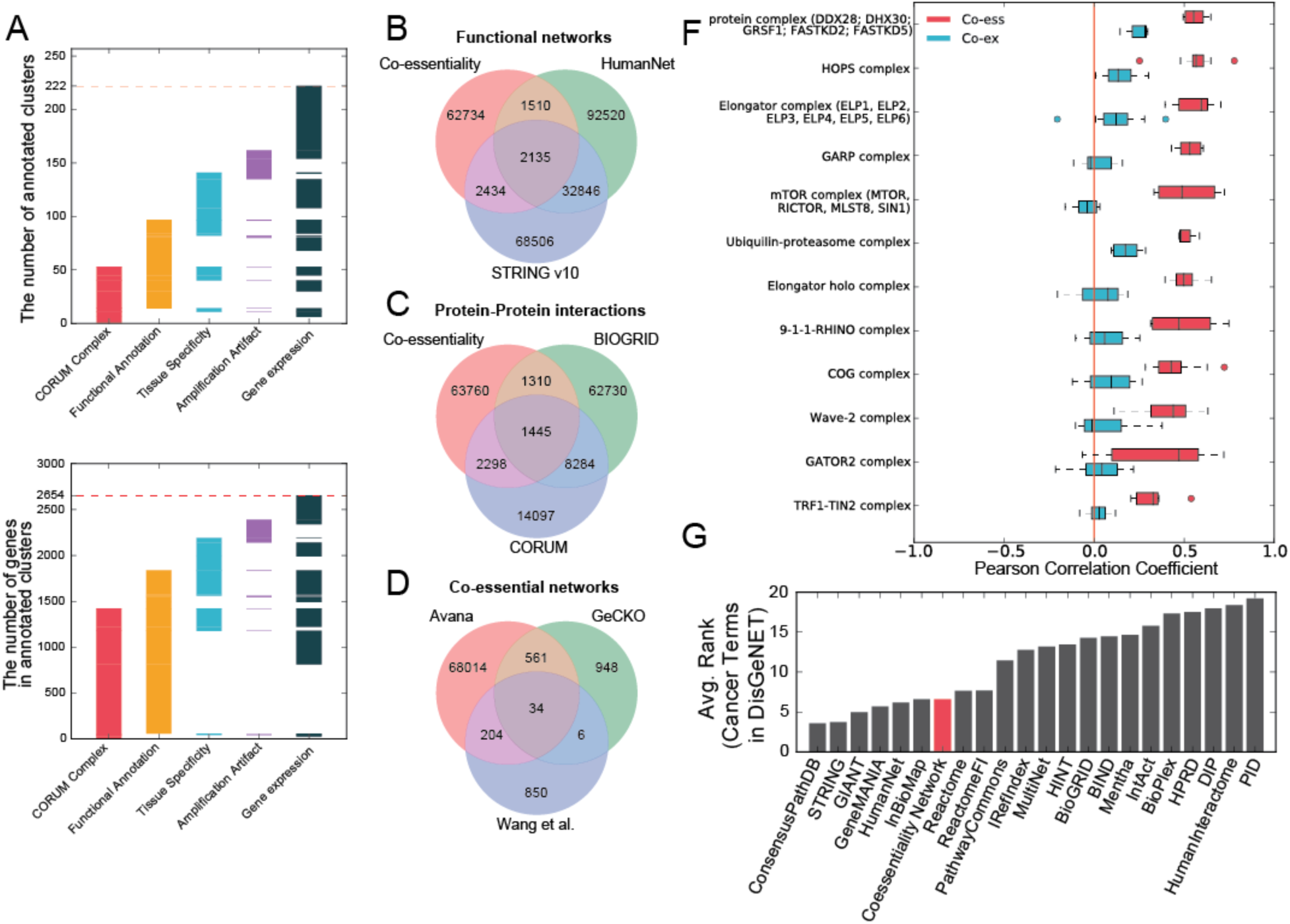
Beyond cancer: characterization of the coessentiality network. (A) Number of clusters (top) and total genes in clusters (bottom) showing strong association with annotated protein complexes, biological function, tissue specificity, amplification-induced CRISPR artifacts, and differential expression of genes. (B,C,D) Comparing the Avana coessentiality network to other functional (B), protein-protein (C), and coessentiality networks (D) shows the unique information contained in our network. (F) For some protein complexes, coessentiality is a better predictor of co-complex membership than co-expression. (G) The coessentiality network is a powerful predictor of cancer pathways (Huang et al.) compared to other databases and networks (lower rank is better).

Given the underlying data, it is perhaps not surprising that oncogenic signatures are clearly evident in the coessentiality network. However, the vast majority of the network structure does not appear to be driven by tissue specificity or mutational signatures. The network contains information complementary to existing functional (Figure 3b) and physical (Figure 3c) interaction networks, and the network derived from Avana data exhibits far greater coverage than equivalent networks from the GeCKOv2 subset of Project Achilles (Aguirre et al., 2016) or Wang (Wang et al., 2017) AML-specific data (Figure 3d). Nevertheless, the remaining network modules show strong functional coherence (Figure 3a). Coessentiality often proves a stronger predictor of complex membership than coexpression (Figure 3e), and this signature is reflected in the network clusters we identified. Indeed 53 clusters, comprised of 1,422 genes, show enrichment for CORUM-annotated protein complexes at P-value < 10^−6^, and fitness profiles have been used to implicate additional members of protein complexes (Pan et al.). However, this holds only for genes whose knockout fitness defects vary across cell lines; coessentiality of core essential genes is poorly predictive of co-complex membership (Supplementary Figure 4). All 53 CORUM-annotated clusters, plus an additional 44 clusters containing 413 genes (totaling 97 network modules with 1,835 genes), show enrichment for GO biological process, cellular component, KEGG pathway, or Reactome pathway annotations at a similarly strict threshold. (All network cluster annotations can be found in the master annotation file, Supplementary Table 7). In addition, we evaluated the relative performance of the coessentiality network by measuring its ability to recover cancer gene sets using DisGeNET (Huang et al., 2018). The coessentiality network ranks comparably with other large functional networks (Figure 3g), while starting from a much smaller data set, suggesting that the coessentiality network explains not only protein complexes but also cancer pathways including interactions between protein complexes and signaling transduction.

Epistatic interactions frequently underlie covariation in fitness profiles (Phillips, 2008). Cluster 2 is highly enriched for genes involved in the mitochondrial electron transport chain, including 30 of 48 genes encoding subunits of NADH dehydrogenase complex (ETC Complex I; P<10^−42^) plus additional subunits of all other ETC complexes. The cluster also contains 49 of 51 subunits of the mitochondrial large ribosomal subunit (P<10^−87^), 23 of 25 members of the small subunit (P<10^−39^), plus 20 mitochondrion-specific tRNA synthases (P<10^−20^). This mitochondrial translation machinery is required for the synthesis of proteins in the ETC complexes. These genes’ inclusion in this cluster, where their essentiality profiles are correlated with those of the complexes they support, reflects a fundamental feature of saturating genetic screens: the essentiality of a given enzyme or biological process is matched by the essentiality of the cellular components required for the biogenesis and maintenance of that process.

We observe numerous additional instances of such epistatic interactions that highlight functional relationships. For example, glutathione peroxidase gene GPX4 shows highly variable essentiality across cell lines (Figure 4a). GPX4 is a selenoprotein that contains the cysteine analog selenocysteine (Sec), the “21^st^ amino acid,” at its active site. Coessential with GPX4 are all the genes required for conversion of serine-conjugated tRNA^Ser^ to selenocysteine-conjugated tRNA^Sec^ (PSTK, SEPHS2, SEPSECS), as well as selenocysteine-specific elongation factor EEFSEC, which guides Sec-tRNA^Sec^ to specific UGA codons. (Figure 4b) (Schoenmakers et al., 2016). Cellular dependence on GPX4 was recently shown to be associated with mesenchymal state (Viswanathan et al., 2017). Our analysis corroborates this finding: we show that GPX4 essentiality is higher in cells expressing mesenchymal marker ZEB1 (P<10^−5^; Figure 4c). In our global analysis, however, GPX4 sensitivity is more highly correlated with the expression level of GPX2, another member of the glutathione peroxidase family (Figure 4d).

**Figure 4.**
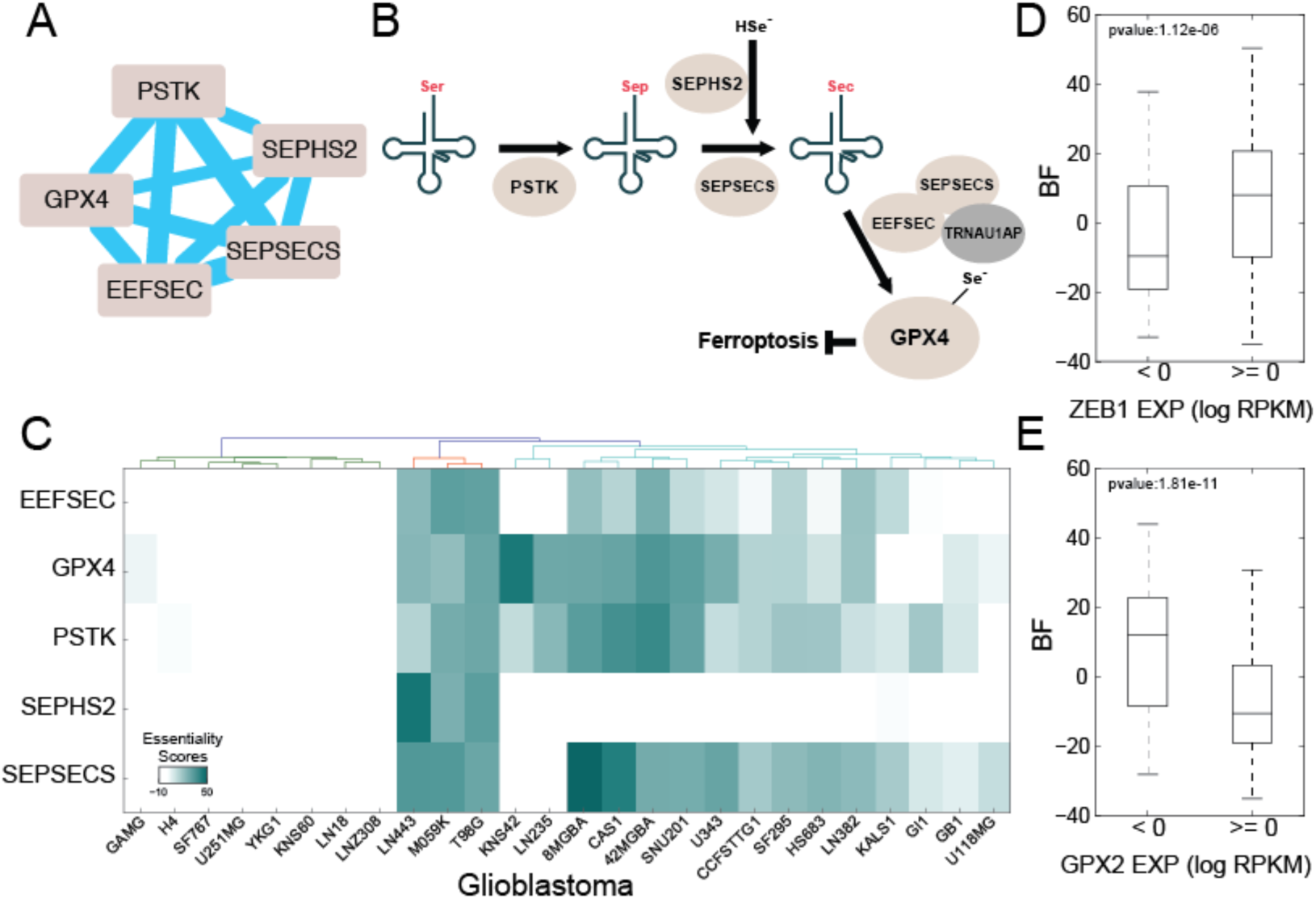
GPX4 cluster. (A) Glutathione peroxidase GPX4, a selenoprotein, is strongly clustered with genes involved in the selenocysteine conversion pathway (B). (C) Entire GPX4 cluster shows marked differential essentiality in glioblastoma cell lines. (D) Cellular requirement for GPX4 is associated with ZEB1 expression, as previously reported, but (E) GPX2 expression is more strongly predictive.

Similarly, a pair of genes, ACOX1 and HSD17B4, which encode three of the four enzymatic steps in peroxisomal fatty acid B-oxidation (FAO), are found in a cluster with ten PEX genes involved in peroxisome biogenesis, maintenance, and membrane transport (Figure 5a-b). The cluster shows a discrete pattern of essentiality, preferentially in lung cells (essential in 6/42 lung cancer lines in the Avana data; Figure 5c) but also appearing intermittently in other lineages. The cluster is not otherwise significantly associated with mutational, copy number, or lineage features. The emergent dependence on peroxisomal FAO is associated with overexpression of 18 genes at P-value < 10^−4^ (See Methods and Supp Table X); two of these include ACOXL, a largely uncharacterized gene similar to ACOX1, and ELOVL7, which catalyzes the rate-limiting step involved in long-chain fatty acid elongation. Notably, this cluster is intact in the network generated from Aguirre et al. (Aguirre et al., 2016)(Figure 5d), though it preferentially arises in pancreatic cells rather than lung cells.

**Figure 5.**
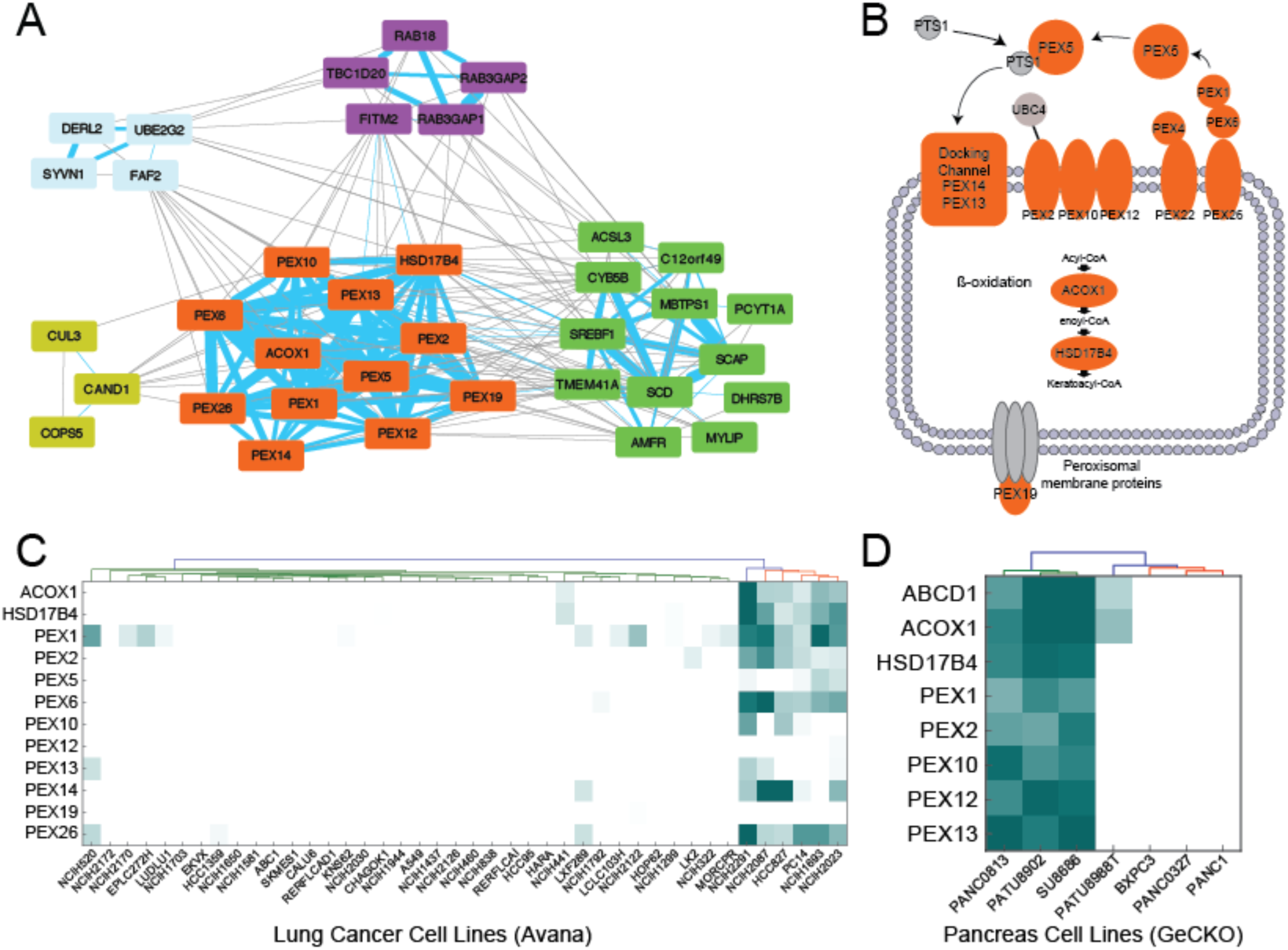
Peroxisomal beta-oxidation. (A) A large group of peroxisome-associated genes (orange) is connected by high-correlation edges in the nework (blue). This cluster is connected by less stringent edges (Benjamini adjusted P-value < 0.01; gray edges) to other clusters containing sterol regulatory genes (green nodes) and the RAB18 GTPase (purple nodes). (B) The PEX cluster contains 12 genes, including 2 enzymes involved in fatty acid oxidation and 10 peroxisome biogenesis and maintenance genes. (C) The PEX cluster is emergently essential in a subset of lung cancer cell lines in the Avana data, and (D) in a subset of pancreatic cancer cell lines in the GeCKO data.

### A network of interactions between biological processes

While individual clusters show high functional coherence, the network of connections between clusters offers a unique window into process-level interactions in human cells. The peroxisomal FAO cluster is strongly connected to another functionally coherent module containing 12 genes, ten of which are tightly connected to other members of the cluster (Figure 5a). Those ten include seven genes whose proteins reside in the ER, five of which regulate cholesterol biosynthesis via posttranslational modification of sterol regulator element binding proteins (SREBPs). The remaining three genes, DHRS7B, TMEM41A, and C12orf49, are largely or completely uncharacterized; their strong association with other genes in this cluster implicates a role in the SREBP maturation pathway. Both the peroxisomal FAO cluster and the SREBP maturation cluster are strongly linked with a module containing RAB18, a RAS-related GTPase involved in Golgi-to-ER retrograde transport, as well as its associated GAP (RAB3GAP1, RAB3GAP2) and GEF (TBC1D120) (Feldmann et al., 2017).

A similar network of modules describes the regulation of the mechanistic Target of Rapamycin (mTOR), in particular its detection of amino acid levels. Figure 6 shows the relationships between a series of network modules describing the core mTOR pathway and several regulatory modules. The mTOR cluster includes mTORC1/2 subunits MTOR, MLST8, MAPKAP1, and RICTOR (mTORC1-specific subunit RAPTOR is never essential and therefore absent from the network); canonical mTORC1/mTORC2 regulatory and signaling components PDPK1, AKT1, and PIK3CB; plus G-protein subunit GNB2, previously shown to physically interact with mTOR in response to serum stimulation (Robles-Molina et al., 2014). Canonical inhibition of mTOR by the TSC1/TSC2 heterodimer – the TSC1-TSC2 link is the top-ranked correlation in the entire dataset, with rho=0.93 (P<10^−117^) – is reflected in the anticorrelation of fitness profiles connecting the TSC1/2 cluster and the mTOR cluster.

**Figure 6.**
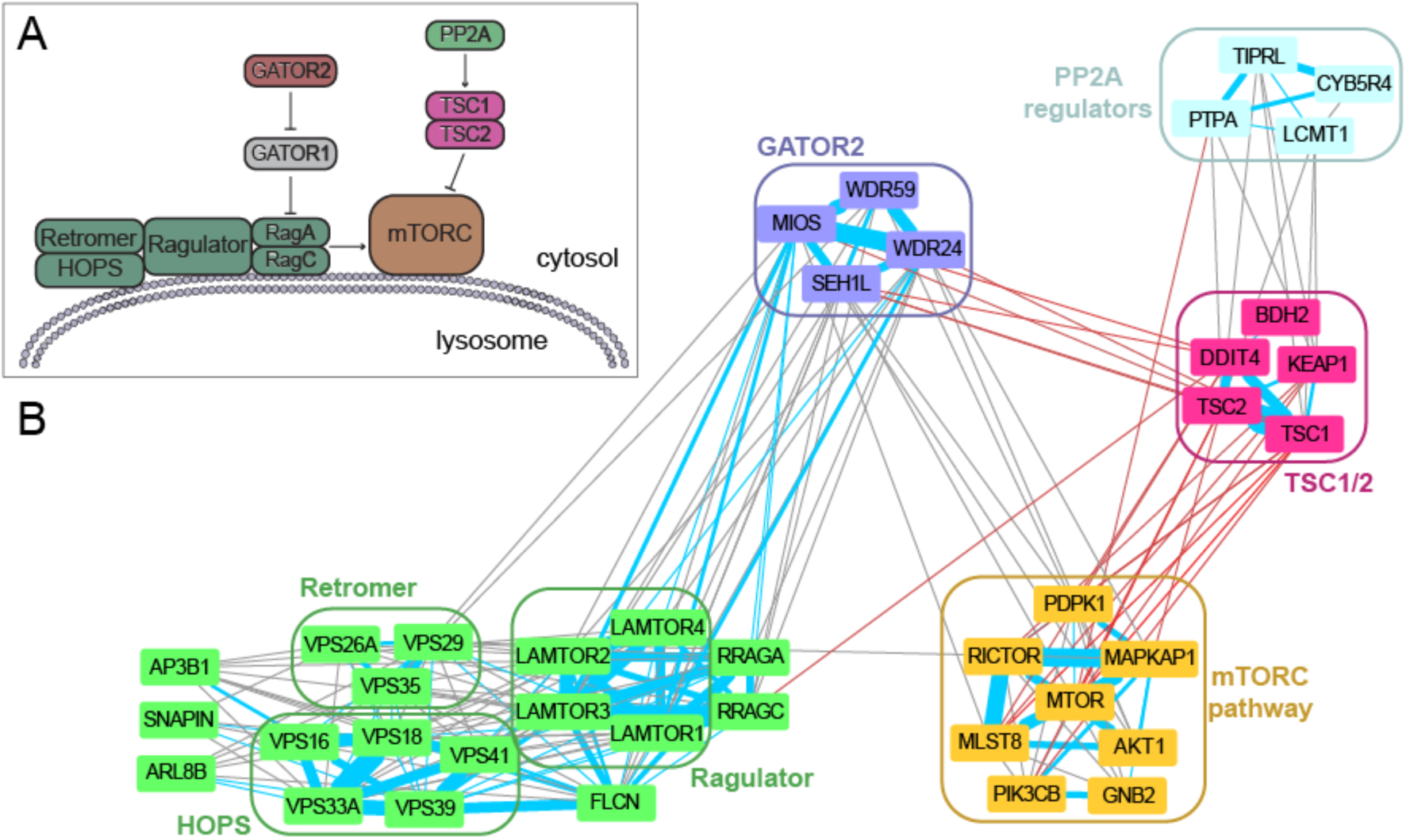
mTORC pathway regulation. (A) The mTORC1/2 complexes are regulated by the canonical TSC1/2 pathway, but amino acid sensing is done via the Ragulator complex at the lysosome. (B) Clusters in the coessentiality network represent components involved in mTORC regulation, and edges between clusters are consistent with information flow through the regulatory network. (Red edges indicate negative correlation).

MTOR response to cellular amino acid levels is modulated by an alternative pathway that functions at the lysosomal membrane (Bar-Peled and Sabatini, 2014). We identify a large cluster containing several genes involved in lysosomal protein and transport, including the HOPS complex (Balderhaar and Ungermann, 2013; Jiang et al., 2014) and the VPS26/29/35 retromer complex (Hierro et al., 2007; Seaman, 2012). This strongly connected cluster also contains the Rag GTPases RagA (RRAGA) and RagC (RRAGC) that transmit information on amino acid abundance to mTORC1 (Bar-Peled and Sabatini, 2014). The Rag GTPases are in turn activated by the Ragulator complex (Bar-Peled et al., 2012; Sancak et al., 2010) and folliculin (FLCN) (Mu et al., 2017), also members of the cluster. The GATOR-1 complex is a nonessential suppressor of essential Rag GTPase activity (Bar-Peled et al., 2013) and is therefore absent from our network, but essential suppression of GATOR-1 by GATOR-2 (Bar-Peled et al., 2013; Wei et al., 2014) is reflected by the strong linkage of the GATOR-2 complex to both the Ragulator and mTOR complexes.

Within the MTOR meta-cluster, we further identify a complex containing three regulators of protein phosphatase 2A (LCMT1, TIPRL, PTPA), whose strong connectivity to the TSC1/2 complex may suggest a regulatory role for PP2A in MTOR signaling. PP2A has previously been posited to be an activator of TSC1/2 upstream of MTOR (Vereshchagina et al., 2008); the coessentiality network suggests specific PP2A regulators that may mediate this regulation.

A third example of the process-level interactions in cells demonstrates the hierarchy of operations required for posttranslational maturation of cell surface receptors. Several clusters in our network describe the ER-associated glycosylation pathways (Figure 7a-b), including synthesis of lipid-linked sugars via the dolichol-phosphate-mannose (DPM) pathway (Ashida et al., 2006; Maeda and Kinoshita, 2008) and extension via the mannosyltransferase family. Glycan chains are transferred to asparagine residues of target proteins via the N-oligosaccharyltransferase (OST) complex. Nascent polypeptide chains are glycosylated as they are cotranslationally translocated into the ER, a process facilitated by signal sequence receptor dimer SSR1/SSR2, and ER-specific Hsp90 chaperone HSP90B1 facilitates proper folding. The OST complex and its functional partners are represented in a single large complex (Figure 7a). Both DPM and OST are highly connected to the large complex encoding GPI anchor synthesis; DPM is required for GPI anchor production (Kinoshita and Inoue, 2000; Watanabe et al., 1998) before transfer to target proteins.

**Figure 7.**
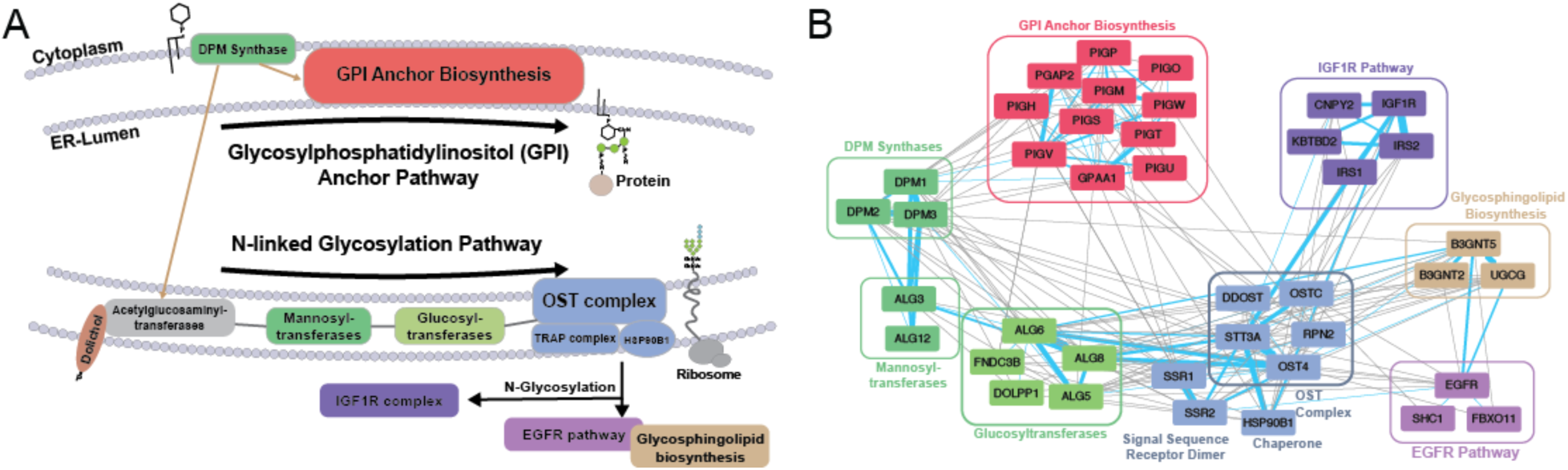
Glycosylation of cell surface receptors. (A) Pathways involved in protein glycosylation and GPI anchor biosynthesis in the ER. (B) A network of clusters around glycosylation tracks the biogenesis and elongation of carbohydrate trees (DPM synthase, mannosyltransferases, glucosyltransferases) to their transfer to target proteins via N-linked glycosylation by the OST complex. Cell surface glycoproteins EGFR and IGF1R are both strongly liked to the OST complex despite their essentiality being mutually exclusive in cell lines.

The variety of oncogenic drivers among the cell lines underlying this network give rise to background-specific dependencies, including a variety of mutated and/or amplified receptor tyrosine kinases (RTKs) with specific, and mutually exclusive, essentiality profiles. Insulin-like growth factor receptor IGF1R is one such RTK, which appears in a cluster with receptor-specific downstream signaling proteins insulin-receptor substrate 1 and 2 (IRS1, IRS2). IGF1R is a highly N-glycosylated RTK and the IGF1R complex is tightly connected to the OST complex in our network. EGFR, also highly glycosylated (Kaszuba et al., 2015), appears in its own cluster with signaling adapter protein SHC1 and is also linked to the OST complex (Figure 7a) despite being mutually exclusive with IGF1R (Supp Figure 2). Interestingly, EGFR is more strongly connected with a separate complex involved in glycosphingolipid biosynthesis (that is itself linked to the OST complex). Prior work suggests that membrane glycolipid composition can strongly influence EGFR autophosphorylation and signaling (Coskun et al., 2011). In contrast, fibroblast growth factor receptor FGFR1 is absent from this meta-network but is strongly associated with heparin sulfate biosynthesis (Supp Fig 2); HS is a known mediator of FGF receptor-ligand interaction (Wu et al., 2003).

## Conclusions

Systematic genetic interaction screens in yeast revealed that most genetic interactions occur either within a biological pathway or between related pathways. We demonstrate that single-gene fitness profiles across screens in genetically diverse human cell lines are analogous to genetic interaction screens across defined isogenic query strains. Importantly, as with model organisms, human genes with correlated fitness profiles are highly likely to participate in the same biological process. We take advantage of this fundamental architectural feature of genetic networks to create a functional interaction map of bioprocesses that demonstrates information flow through a human cell. The network predicts gene function and provides a view of process-level interactions in human cells, allowing a level of abstraction beyond the gene-centric approach frequently employed.

The network is derived from the emergent essentiality of defined biological processes and the genes required to execute them. We show that this approach significantly expands our knowledge beyond current networks of comparable design (e.g. STRING, HumanNet). A critical next step will be to understand the underlying context that drives the emergent essentiality of specific bioprocesses in specific backgrounds. The health implications of this question are profound. In cancer, to understand the causal basis of modular emergent essentiality is to identify matched pairs of biomarkers (the causal basis) and precision targets (the essential pathway) for personalized chemotherapeutic treatment. Additionally, lineage-specific essential processes could provide explanatory power for germline mutations causing tissue-specific disease presentation, in cancer as well as other diseases.

Expanding the coverage of the network will require different screening approaches. Fitness screens in cancer cell lines in rich media will miss cellular dependencies that are present only under stress conditions. In yeast (Hillenmeyer et al., 2008) and nematodes (Ramani et al., 2012), these context-dependent fitness effects comprise the majority of genes in the genome. Increasing the coverage of the genetic interaction network beyond the ∼3,000 genes whose fitness profiles covary across human cancer cell lines will require screening in different nutrients and perturbagens, as well as sampling the effects of genetic mutations outside common cancer genotypes. Nevertheless, the indirect approach to identifying genetic interactions from monogenic perturbation studies is demonstrably effective, and offers a powerful tool for navigating the network of connections between cellular bioprocesses. The coessentiality network used in this study can be viewed interactively at https://pickles-hartlab.shinyapps.io/cyto_app/ and downloaded at the NDEx project.

## Methods

See Supplementary Data for a complete description of methods used in this study.

## Acknowledgments

The authors would like to acknowledge the scientists, administrators, and funding agents behind Project Achilles. Without their commitment to rapid release of open access data, none of this work would have been possible.

Funding was supplied by the MD Anderson Cancer Center Support Grant P30 CA016672 (the Bioinformatics Shared Resource) and the Cancer Prevention Research Institute of Texas (CPRIT) grant RR160032.

